# Beta-band activity in medial prefrontal cortex predicts source memory encoding and retrieval accuracy

**DOI:** 10.1101/437244

**Authors:** Karuna Subramaniam, Leighton B.N. Hinkley, Danielle Mizuiri, Hardik Kothare, Coleman Garrett, Anne Findlay, John F. Houde, Srikantan S. Nagarajan

## Abstract

Reality monitoring is defined as the ability to distinguish internally self-generated information from externally-derived information. The medial prefrontal cortex (mPFC) is a key brain region subserving reality monitoring and has been shown to be activated specifically during the retrieval of self-generated information. However, it is unclear if mPFC is activated during the encoding of self-generated information into memory. If so, it is important to understand whether successful retrieval of self-generated information critically depends on enhanced neural activity within mPFC during initial encoding of this self-generated information.

We used magnetoencephalographic imaging (MEGI) to determine the timing and location of cortical activity during a reality-monitoring task involving self generated contextual source memory encoding and retrieval. We found both during encoding and retrieval of self-generated information, when compared to externally-derived information, mPFC showed significant task induced oscillatory power modulation in the beta-band. During initial encoding of self-generated information, greater mPFC beta-band power reductions occurred within a time window of −700ms to −500ms prior to vocalization, activity in mPFC that was not observed during encoding of externally-derived information. This mPFC activity during encoding of self-generated information predicted subsequent retrieval accuracy of self-generated information. Beta-band activity in mPFC was also observed during the initial retrieval of self-generated information within a time window of 300 to 500ms following stimulus onset and correlated with accurate retrieval performance of self-generated information. Together, these results further highlight the importance of mPFC in mediating the initial generation and awareness of participants’ internal thoughts.

## Introduction

Reality monitoring is defined as the ability to distinguish the source of internally self-generated information from outside reality (externally-derived information)^1–4^. The medial prefrontal cortex (mPFC) is a key brain region that mediates reality monitoring and has also been shown to be activated during the retrieval of self-generated information. Although most studies have found mPFC activity increase during the retrieval of self-generated information^3–8^, to-date no study has investigated whether mPFC is activated during the encoding of self-generated information on the same task. If so, it is important to delineate the timing in the oscillatory frequency band power of mPFC activation to investigate when mPFC activity during encoding of self-generated information may be able to predict subsequent retrieval of this information.

In this study, we use magnetoencephalography (MEG) imaging to delineate the neural basis of the initial encoding of one’s own thoughts into memory prior to the actual retrieval/identification of these self-generated thoughts. In particular, we use MEG to examine the timing of precisely when induced neural oscillatory activity mediating the initial encoding of self-generated information can predict and facilitate accurate retrieval of this self-generated information. During the encoding of self-generated information, participants need to generate their own internal thoughts and then transform these thoughts into actions by vocalizing their internally generated word during a sentence completion task, as opposed to the externally-derived condition in which the final word is already provided by the experimenter. During both encoding of self-generated and externally-derived information, participants engage in the same goal-directed action of vocalizing the final word on matched semantically constrained sentences (**Fig. 1**). Therefore any neural differences that result between the self-generated versus externally-derived conditions must be attributable to neural correlates specifically mediating the generation of one’s own internal thoughts and the cognitive preparation and volition to initiate and transform these internally-generated thoughts into self-initiated actions (speech execution). During accurate retrieval of self-generated information, we have previously consistently found increased activation within the medial prefrontal cortex (mPFC) when compared to retrieval of externally-derived information^3–6^. The primary focus of the present study is to examine whether successful identification of self-generated information critically depends on neural activity during encoding of this self-generated information in mPFC-enhanced activity.

**Figure 1.**
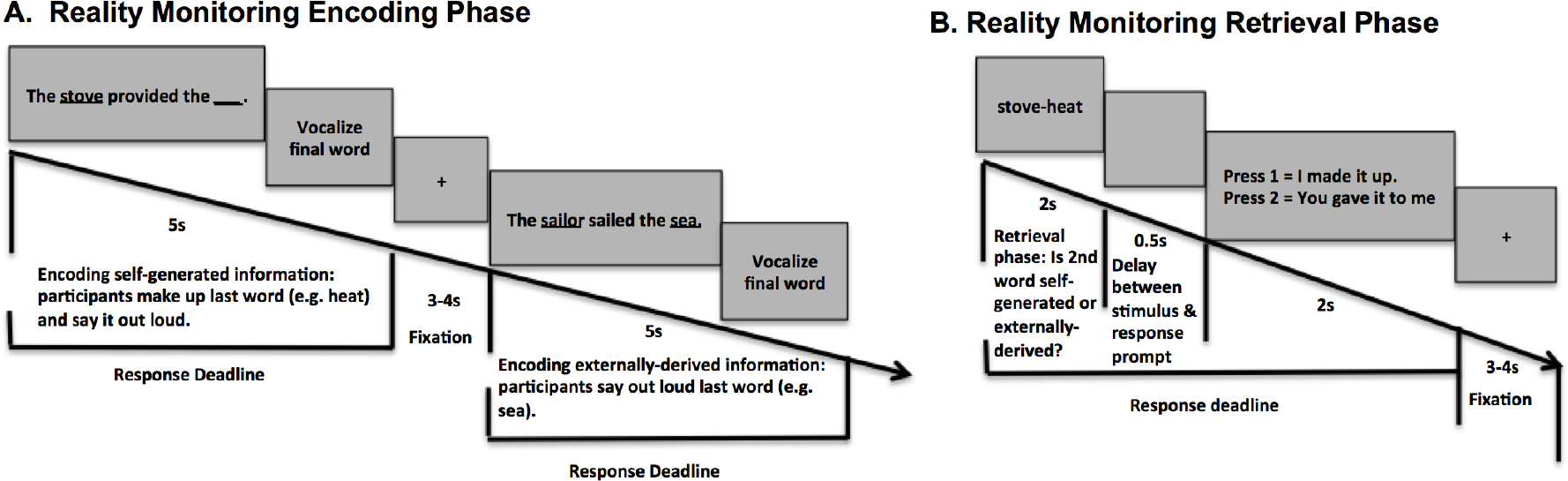
Reality Monitoring Task Design. A. Encoding Task. Participants were given sentences in which the final word was either left blank for participants to generate and vocalize themselves (e.g., *The <u>stove</u> provided the* _) or was externally-derived as it was provided by the experimenter for subjects to vocalize (e.g., *The <u>sailor</u> sailed the <u>sea</u>*). **B. Retrieval Task:** Participants were randomly presented with the noun pairs from the sentences (e.g., *stove-heat*), and had to identify with a button-press whether the second word was previously self-generated or externally-derived.

Reconstructions of MEG sensor data using adaptive spatial filtering techniques^9^ provides precise source localizations during encoding and retrieval of self-generated information, allowing us to examine both the timing and location of neural activity from internal thought generation to preparation and volition to transform these intentions into actions. A recent study has shown that examining changes in beta oscillations (12-30Hz) is a robust way of examining neural differences between self-generated versus externally-derived changes prior to an action^10^. Beta band suppression has additionally been used as a reliable marker of neural activity increases that precede self-initiated actions of one’s own volition^10–12^. However, it is unknown whether such beta band suppression that precedes self-generated actions can be observed in mPFC – specifically during the encoding of self-generated information and, if so, whether this activity can predict subsequent successful retrieval of this self-generated information. If we find that mPFC is activated during the encoding of self-generated information, MEG source analyses will allow us to pinpoint the timing of activation in mPFC and probe whether mPFC is activated at a specific time during the initial generation of one’s own thoughts (but not during the externally-derived condition) prior to motor preparation and execution.

To summarize, we examine the following three hypotheses: 1) During encoding of self-generated information when compared to externally-derived information, increased beta suppression (indexing enhanced neural activity) would be observed in mPFC. 2) During retrieval of self-generated information when compared to externally-derived information, we would find increased beta suppression (indexing enhanced neural activity) in mPFC. 3) Activity in mPFC during both encoding and retrieval of self-generated information would be associated with accurate retrieval performance. Support for these hypotheses will not only indicate that mPFC mediates the successful encoding of self-generated information, but also that its activity predicts and facilitates subsequent retrieval and accurate identification of this self-generated information.

## Materials and Methods

### Participants

In the present study, we recruited 15 healthy participants who completed the reality monitoring task (7 females, mean age = 31.20, range = 21-50, mean education = 19.03). This study was approved by the Internal Review Board (IRB) at the University of California San Francisco (UCSF) and all research was performed in accordance with IRB regulations at UCSF. All participants gave written informed consent and then completed the reality monitoring task. Inclusion criteria for healthy participants were: no psychiatric/neurological disorders, no substance dependence or current substance abuse, meets MRI criteria, good general physical health, age between 18 and 60 years, right-handed and English as first language.

### Reality Monitoring Task

All subjects completed the reality monitoring task in the MEG scanner. Reality monitoring requires that subjects make higher-order conscious judgments about distinguishing whether information was previously self-generated or externally-derived. As described in previous experiments, the reality monitoring task consisted of an encoding phase and a memory retrieval phase conducted in separate runs^4^ (see **Fig. 1A,B**). During encoding, at the beginning of every trial, participants were visually presented with 200 semantically constrained sentences with the structure “noun-verb-noun,” presented in blocks of 20 trials per run. On alternating half of the sentences, the final word was either left blank for participants to generate themselves (e.g., *The <u>stove</u> provided the* **__**) or was externally-derived as it was provided by the experimenter (e.g., *The <u>sailor</u> sailed the <u>sea</u>*). For each sentence, participants were told to pay attention to the underlined words and to vocalize the final word of each sentence (**Fig. 1A**). Each trial was 5s in length, with a variable fixation period between 3-4s. In a separate run, participants completed the reality-monitoring retrieval task where they were randomly presented with the noun pairs from the sentences (e.g., *stove-heat*) at the beginning of every trial for a 2s period, and had to identify with a button-press whether the second word was previously self-generated or externally-derived (**Fig. 1B**). The number of correctly identified self-generated and externally-derived trials was computed for each participant during the retrieval phase.

### Behavioral Statistical Analyses

Repeated-measures ANOVAs were implemented in SPSS to examine differences in accuracy and reaction times (RT) between correctly identified self-generated and externally-derived information. Outliers were defined as values above/below 2 standard deviations from the mean. We did not find any outliers in the behavioral data. Mean accuracy and RT were computed for correctly identified self-generated and externally-derived information, averaged across all participants (**Table 1**).

**Table 1.** Reality-monitoring Accuracy and Reaction Times (RT)

|  | **Self-generated**<br>**Identification** | **Externally-derived**<br>**Identification** | **Self vs. External**<br>**Identification (p**<br>**value)** |
| --- | --- | --- | --- |
| **Accuracy (%)** | 83.83 ± 9.77 | 83.13 ± 9.13 | p = 0.82 |
| **RT (ms)** | 1212.83 ± 124 | 1359.16 ± 156 | p < 0.0001 |

### MEG Data Acquisition

Magnetic fields were recorded in a shielded room using a whole-head 275 axial gradiometer MEG system with third-order gradient correction (MEG International Services Ltd. (MISL), Coquitlam, BC, Canada) at a sampling rate of 1200 Hz and acquired under a bandpass filter of 0.001–300 Hz. Three fiducial coils (nasion, left/right preauricular) were placed to localize the position of the head relative to the sensor array in order to be later co-registered onto the structural MRI to generate head shape. Head localization was performed at the beginning and ending of each task block to register head position and to measure head movement during the task. Third gradient noise correction filters were applied to the data and corrected for a direct-current-offset based on the whole trial. Noisy sensors and trials with artifacts (i.e., due to head movement, eye blinks, saccades or sensor noise) were defined as magnetic flux exceeding 2.5 pT. Epochs were rejected from further analysis if they contained artifacts. High-resolution structural MRI images were also acquired sagittally on a 3-Tesla Siemens MRI scanner at the Neuroscience Imaging Center (MPRAGE sequence; 160 1mm slices, FOV=256mm, TR=2300ms, TE=2.98ms) in order to reconstruct MEG data in source space.

### MEG Data Analyses

Spatiotemporal estimates of neural sources were generated using a time–frequency optimized adaptive spatial filtering technique implemented in the Neurodynamic Utility Toolbox for MEG (NUTMEG; http://nutmeg.berkeley.edu) to localize induced changes in oscillatory power, with focus on beta (12-30Hz) band. Adaptive spatial filtering was applied within the beta band frequency which also provides additional robustness to eye-blink and saccade artifacts^13,14^. A tomographic volume of source locations (voxels) was computed through an adaptive spatial filter (5 mm lead field) that weights each location relative to the signal of the MEG sensors^9^. Source power for each location was derived through a noise-corrected pseudo-F statistic expressed in logarithmic units (decibels) comparing signal magnitude during an “active” experimental time window versus a baseline “control” window^15^. Datasets were independently reconfigured into stimulus (0ms = visual stimulus set presentation) and response-locked (0ms = vocal or button press onset, respectively) formats for separate analyses during the encoding and retrieval phases of the reality-monitoring task. Sliding window lengths in beta band (with 200ms window size) and sampling rate were examined for activity throughout the time-course of each task condition during the stimulus-locked and response-locked periods, and were compared to a resting fixation (inter-trial) baseline window of the same length as the active window.

We focus here on source-space reconstructions in the beta (12–30 Hz) band given that suppression in this frequency range is related to cortical activation^16–19^ and has been found to be a reliable marker of neural activity increases that precede self-initiated actions^10^. High-resolution anatomical MRIs were spatially normalized to a standard MNI template using SPM8 (http://www.fil.ion.ucl.ac.uk/spm/software/spm2/) with the resulting parameters being applied to each individual subject’s source-space reconstruction through NUTMEG. Group analyses to evaluate effects at the second level were performed with statistical nonparametric mapping^20^. To minimize spatial frequency noise in the beamformer volumes, average and variance maps for each individual time window were calculated and smoothed using a Gaussian kernel with a width of 20×20×20 mm FWHM^21^. From these volumes, a pseudo-F statistic is obtained for each voxel, time window, and frequency band. Statistical significance was estimated by obtaining a permuted distribution (through 2^N^ possible combinations of negations) and estimating the significance of each pseudo-F value from its position in this permuted distribution^20^. In order to examine activation specifically induced by self-mediated processes, we contrasted whole-brain neural activity during the encoding and retrieval of self-generated information with encoding and retrieval of externally-derived information by subtracting these two periods and running a one-sample group t-test on the subtraction maps^9^. Additionally, we computed whole-brain correlations in NUTMEG (Pearsons’s r) between induced neural oscillatory activity during encoding and retrieval of self-generated information with behavioral measures of performance (i.e., accurate identification of this self-generated information). Effect sizes (Cohen’s d) were used to quantify the power of the linear relationships. We controlled for confounds from behavioral performance by examining neural activity mediating only correct performance during the retrieval phase of the reality monitoring task. For all analyses, multiple-comparisons corrections of the uncorrected thresholds at p<0.001 derived from non-parametric tests were then applied using an additional cluster correction (at least 50 contiguous voxels).

## Results

### Behavioral Results

Mean accuracy and RT data during retrieval of self-generated and externally-derived information in the reality monitoring task is shown in **Table 1**. Participants were faster at identifying self-generated information when compared to externally-derived information (F[1,14]=27.51, p<.0001) but were not more accurate (F[1,14]=0.05, p=.82) (see Table 1). We did not find any difference in the number of noisy trials that were removed due to movement, eye-blinks or saccades between self-generated and externally-derived trials during either encoding (F[1,14]=1.00, p=.33) or retrieval (F[1,14]=.09, p=.76).

### MEG Results

During the encoding phase, we examined whole-brain fluctuations in beta (12–30 Hz) oscillatory activity in the period after stimulus onset (at presentation of the sentence) and prior to vocalization onset of the final word (0 ms). We found that participants showed greatest activation (p<.001, corrected) in mPFC during encoding of self-generated information (i.e., during the contrast of self-generated vs. externally-derived information), between −700 to −500ms prior to vocalization onset (**Fig. 2**). Prior to −700ms before vocalization onset, we did not find any mPFC activity, providing evidence that mPFC activity cannot be attributed to early cognitive processing differences between the two types of stimuli during initial stimuli presentation (**Fig. 2**). The lack of mPFC activity closer to vocalization onset (between −500ms to 0ms) additionally demonstrates that mPFC activity does not appear to mediate a general motor preparation to act (i.e., vocalize one’s thoughts). These findings suggests that mPFC was only active within a specific time window prior to this point, which likely represents the initial generation of one’s own internal thoughts, and the cognitive volition to transform these internally-generated thoughts into actions (speech output).

**Figure 2.**
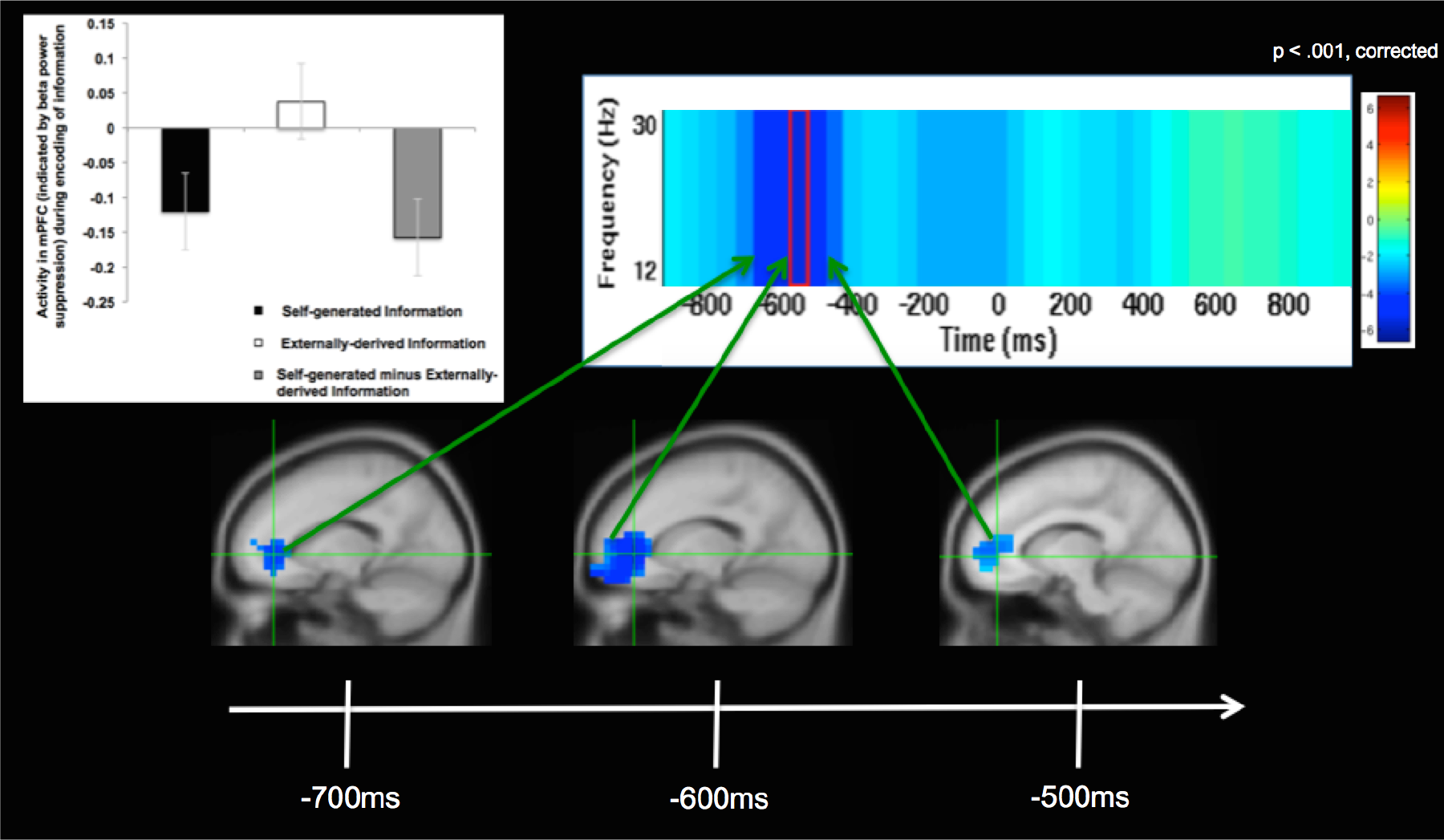
Self-generated Information Encoding Time Windows. Response-locked (0ms=vocalization onset) group analyses of changes in beta (12-30Hz) oscillatory power, indicating significant increases in mPFC activity during self-generated encoding (i.e., during encoding of self-generated vs. externally-derived information) only within the specific limited time windows shown between −700 to −500ms prior to vocalization onset. All images are cluster corrected (>50 contiguous voxels) on statistical maps thresholded at p < 0.001, cross-hairs centered at the peak of mPFC activity (x,y,z = −15,40,0). Bar charts illustrate beta weights averaged across all participants for each condition, and are shown at the peak of mPFC activity during encoding of self-generated vs. externally-derived information.

During the retrieval phase, we also examined whole-brain fluctuations in beta oscillatory activity in the period after stimulus onset at presentation of the word-pair (0 ms) up to the button-press (i.e., when subjects identified whether the final word of the sentence was previously self-generated or externally-derived). We only examined trials with accurate identification. Our whole-brain analyses revealed that participants not only showed greatest activation (p<0.001, corrected) in mPFC during accurate retrieval of self-generated information (i.e., during the contrast of self-generated vs. externally-derived information) within a specific time window between 300ms to 500ms after stimuli word-pair onset (**Fig. 3**) but also that participants also showed the same peak in mPFC activity during encoding of self-generated information that they showed during accurate retrieval of this self-generated information (i.e., same peak co-ordinates; x=−15, y=40, z=0) (**Figs. 2 & 3**). Furthermore, the lack of mPFC activity immediately prior to the button-press (i.e., after 500ms) negates the idea that mPFC activity could be attributed to a general motor preparation to act (**Fig. 3**). Rather, these findings indicate that mPFC activity observed during these specific time windows likely represents the initial generation and awareness of one’s own thoughts and the volition to cognitively prepare to transform these internal thoughts into self-initiated actions.

**Figure 3.**
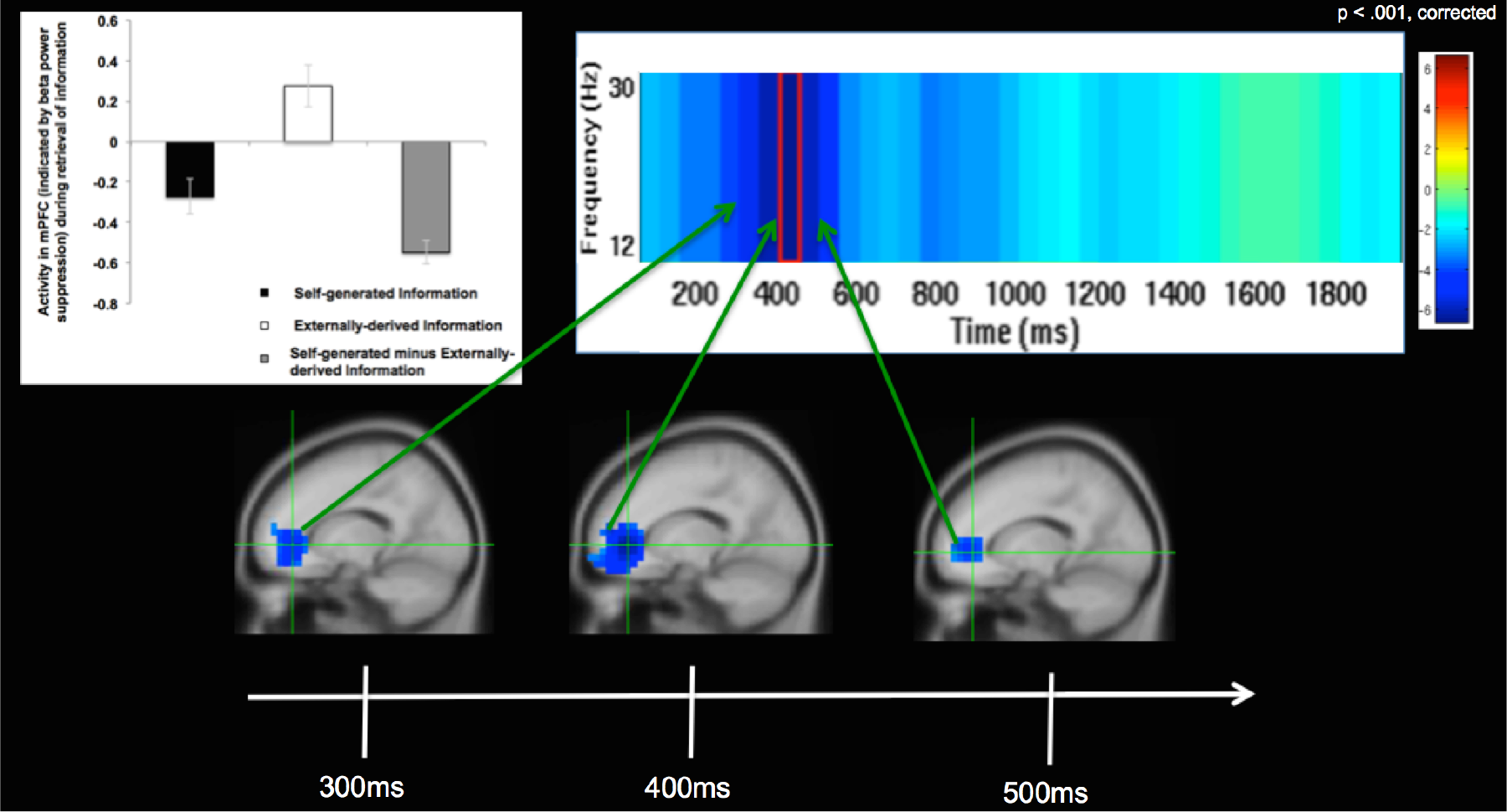
Self-generated Information Retrieval Time Windows. Stimulus-locked (0ms=word-pair onset) group analyses of changes in beta (12-30Hz) oscillatory power, indicating significant increases in mPFC activity during self-generated retrieval (i.e., during retrieval of self-generated vs. externally-derived information) only within the specific limited time windows shown between 300 to 500ms after stimulus onset. All images are cluster corrected (>50 contiguous voxels) on statistical maps thresholded at p < 0.001, cross-hairs centered at the peak of mPFC activity (x,y,z = −15,40,0). Bar charts illustrate beta weights averaged across all participants for each condition, and are shown at the peak of mPFC activity during retrieval of self-generated vs. externally-derived information.

We also performed whole brain-behavior correlation analyses between neural activity (12-30Hz) observed during the encoding and retrieval of self-generated information with participants’ accurate identification of this information as self-generated. These whole-brain correlation analyses also resulted in large significant (p<0.001, corrected) mPFC clusters **(Figs. 4 & 5)**, which overlapped with the mPFC cluster observed during encoding of self-generated information (i.e., during the contrast of self-generated vs. externally-derived information) within these same time windows (i.e., between −700 to −500ms prior to vocalization for encoding; between 300-500ms after stimuli onset for retrieval). Cohen’s d analyses yielded large effect sizes for the correlation coefficients between mPFC activation during encoding and retrieval of self-generated information with behavioral accurate identification of this self-generated information (Cohen’s d = 1.5 and 2.14, respectively). These findings indicate that enhanced mPFC activity during encoding of self-generated information (indicated by increased suppression of the beta (12-30Hz) rhythm over mPFC) primed and potentiated subsequent retrieval and accurate identification of this self-generated information.

**Figure 4.**
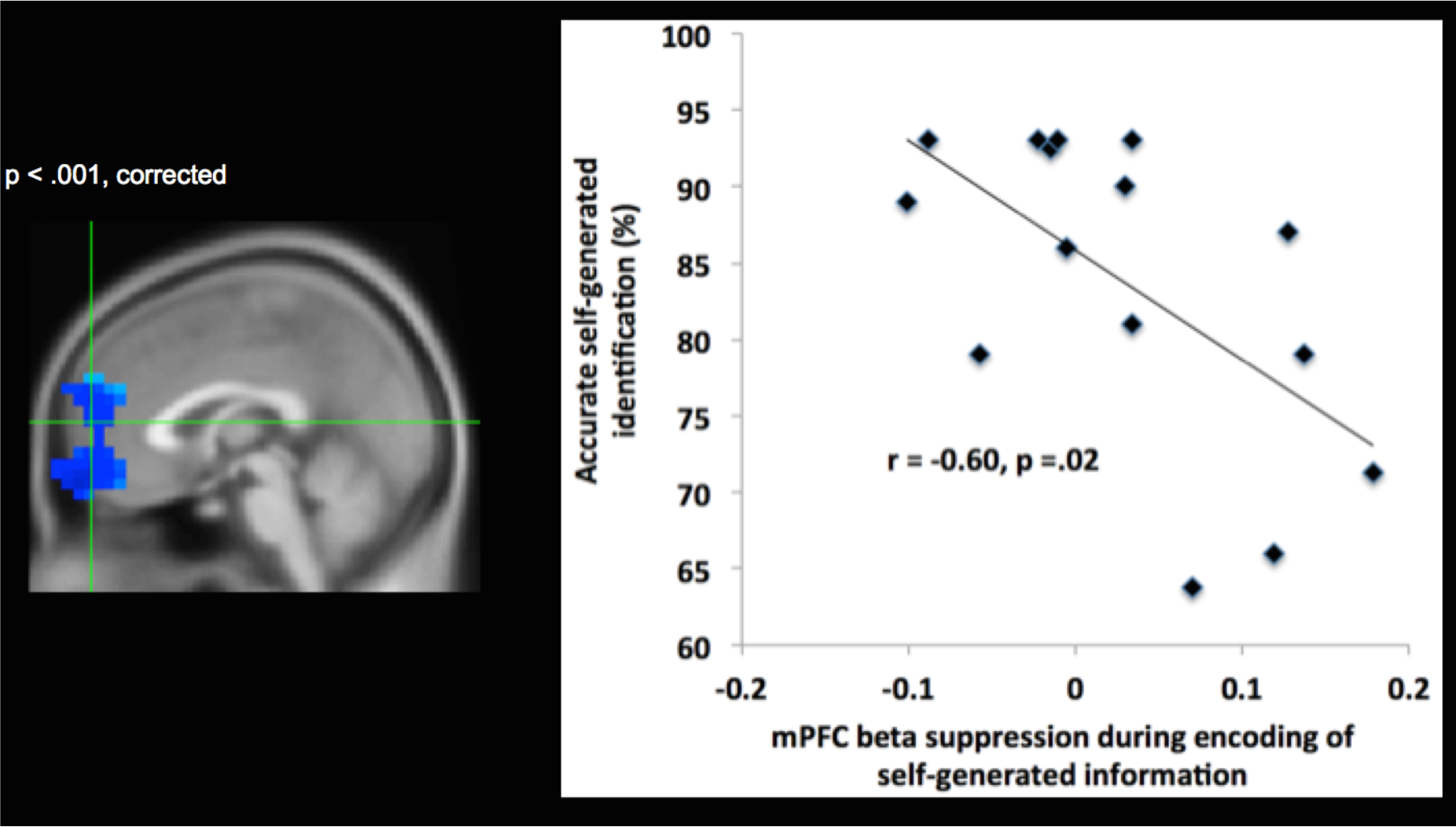
Brain-behavior correlation during encoding of self-generated information. Whole-brain correlation analyses, indicating that activity in mPFC (shown by changes in beta (12-30Hz) oscillatory power) during encoding of self-generated information significantly correlates with subsequent behavioral accurate identification of this self-generated information only within specific time windows between −700 to −500ms prior to vocalization onset (thresholded at p <0.001 and cluster corrected at >50 voxels, peak brain-behavior correlation activity at x,y,z = 1,60,−17).

**Figure 5.**
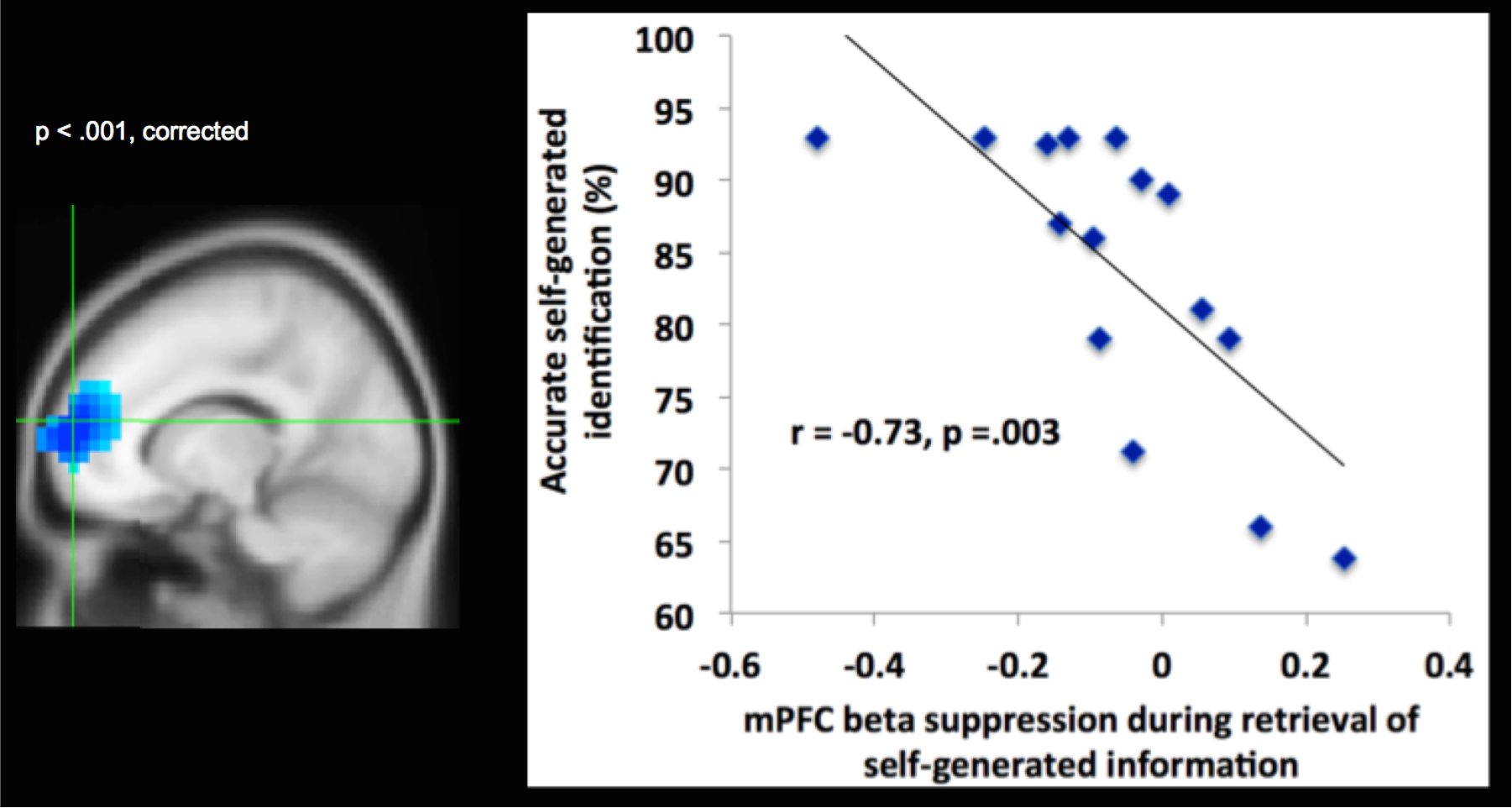
Brain-behavior correlation during retrieval of self-generated information. Whole-brain correlation analyses, indicating that activity in mPFC (shown by changes in beta (12-30Hz) oscillatory power) during retrieval of self-generated information significantly correlates with behavioral accurate identification of self-generated information within time windows between 300 to 500ms after stimulus onset (thresholded at p < 0.001 and cluster corrected at >50 voxels, peak brain-behavior correlation activity at x,y,z = −15,60,10).

Importantly, we found that mPFC was significantly activated (indicated by beta power suppression) only for the self-generated condition during both encoding and retrieval of information, which correlated with accurate self-generated identification. Note that during both encoding and retrieval of externally-derived information, mPFC was deactivated (indicated by beta power increases above baseline). Therefore, this mPFC activation that occurred only for the self-generated condition was the driving force behind the highly significant difference between self-generated and externally-derived information, as illustrated by the bar charts (**Figs. 2 & 3**).

## Discussion

We provide convergent evidence from different analyses that mPFC is significantly activated for the self-generated condition within specific time windows during both encoding and retrieval of information. The same locus of mPFC peak activity was found during both encoding and retrieval of self-generated information, indicating that the same region within mPFC appears to mediate the initial generation and awareness of one’s own thoughts, and correlates with subsequent accurate identification of these self-generated thoughts and actions. Task induced mPFC activity was specific to the encoding and retrieval of self-generated information (i.e., mPFC was not activated during externally-derived information), which resulted in the highly significant difference between self-generated and externally-derived information. Importantly, mPFC activation was found within a limited specific time window during both encoding and retrieval of self-generated information; this activation occurred prior to motor preparation and execution during which time people were likely to have first generated and become aware of their internally-generated thoughts. Together, these convergent findings indicate that mPFC likely represents an early neural correlate of the ‘self’ that mediates the initial generation and awareness of one’s own thoughts and the volition to cognitively prepare to transform these self-generated thoughts into actions.

On our reality-monitoring task, the self-generated condition requires a greater sense of volition and agency in the preparation and generation of internally self-generated thoughts and their transformation into actions because participants need to decide what word to generate, as opposed to the externally-derived condition in which the final word is already provided by the experimenter. These findings are consistent with prior research suggesting that the prefrontal cortex may be one candidate that mediates the volition to transform one’s own internally-generated intentions to actions^23^, and likely represents an early neural correlate of self-agency. Self-agency is defined as the experience of being the agent of one’s own thoughts and actions^24–26^. Self-agency depends on successful encoding and memory retrieval of one’s own thoughts and actions, thus enabling accurate judgments that ‘I generated my own thoughts and actions’^26^. Our findings demonstrating that mPFC activity increase was significant only within specific time windows during encoding and retrieval, indicate that mPFC activity likely reflects subjects’ initial awareness of being the agent of their own thoughts. For example, the lack of mPFC activity observed yoked to stimuli sentence onset during encoding indicates that mPFC activity was not due to early cognitive processing differences between the two types of stimuli during sentence presentation. The lack of mPFC activity immediately prior to response onset also suggests that mPFC activity does not also simply mediate a general motor preparation to act, but seems to be specific to the initial generation and volition to transform these internal thoughts to self-initiated actions in order to produce the resulting experience of self-agency. This also explains why we did not observe mPFC activation during the encoding of externally-derived words during this period (**Fig. 2**).

During retrieval of self-generated information, mPFC activity increase was significant only within a specific time window observed between 300-500ms after stimulus onset which likely reflects subjects’ initial awareness of being the agent of their own thoughts (i.e., the awareness that information is previously self-generated). We also note here that because mPFC activity increase during this specific time window occurred well before the actual button-press (i.e., which occurred roughly between 1200-1400ms across subjects), mPFC increase in activity cannot be attributable to a general motor preparation to act or to any differences in performance on the task (i.e., RT and accuracy), but rather likely mediates the initial awareness of self-agency. We think it is also unlikely that mPFC activity during the self-generated condition can be attributable to increased working memory load or effortful processing. If this was true, since participants took longer to identify externally-derived information (likely indicating more cognitive effort), we would expect mPFC activity to be greater during the identification of externally-derived information, rather than what we find here for self-generated information. Finally, because we also observed significant mPFC activation during the retrieval of self-generated information and mPFC deactivation during externally-derived information retrieval, we can conclude that mPFC activity is specific to self-generated condition, and likely mediates the initial awareness of the experience of self-agency.

Our findings are consistent with prior research indicating that the mPFC appears to be particularly implicated in tagging information as being relevant to the “self” ^3,27–29^. For example, Cabeza et al.^7^ found that mPFC activation was greater when subjects viewed photographs that they themselves had taken (the autobiographical “self” condition) versus when they viewed similar photographs taken by another person (the “external” condition), and that this enhanced mPFC activation difference was driven by mPFC activity increase specifically during the “self” condition. Our study was hypothesis driven to examine beta-band activity in mPFC, particularly in light of our and other prior studies which have demonstrated a consistent preparatory process prior to self-generated actions, indexed by EEG beta band power reductions termed as a “readiness potential” which does not occur before externally-derived thoughts and actions but is specific to the planning, preparation and initiation of self-generated thoughts and actions^10,30,31^. In this recent study, Khalighinejad et al^10^ demonstrated that such self-initiated actions were preceded by a reduction in noise variability in EEG beta band power reductions, potentially indicating a neural precursor of self-initiated actions. For the first time, here, we provide evidence that the mPFC may be one candidate that represents one underlying neural correlate mediating this readiness potential. This is because not only is mPFC activated prior to self-initiated actions and initial awareness of self-agency during retrieval, but more importantly because it is also preferentially activated during the initial generation and encoding of one’s internal self-generated thoughts and correlates with subsequent accurate retrieval performance, indexing an encoding precursor to early self-awareness and self-agency. A more comprehensive data-driven approach to examine other frequency bands in a larger oscillatory network is warranted in the future.

## Conclusion

In conclusion, we show that the mPFC mediates initial encoding and early awareness of one’s internal self-generated thoughts. We show that mPFC activity increase was not simply due to a relative increase in activity that resulted from the difference between self-generated and externally-derived conditions, but was specifically related to the self-generated condition. Importantly, mPFC activity increase was found within a specific limited time window that occurred during the encoding and early awareness of one’s own thoughts, and its activity correlated with subsequent accurate judgments of self-agency within these same time windows. Taken together, the present findings demonstrate that mPFC is a good candidate for representing one neural precursor or early neural correlate of the experience of self-agency, which is activated during the initial generation and awareness of internally self-generated thoughts and actions.

## Acknowledgments

This research was supported by the Brain and Behavior Research Foundation Young Investigator Award grant (NARSAD: 17680) and NIMH K01 grant (KO1MH82818) to Karuna Subramaniam, and the following NIH grants to Srikantan Nagarajan and John Houde (R01DC010145, R01DC013979, and R01NS100440).

## Additional Information

### Author Contributions

KS recruited all the subjects and designed the reality monitoring experiment; acquired, analyzed the reality monitoring data and interpreted the data; and wrote and edited the manuscript. LH helped with acquisition and analyses of the MEGI data. DM helped with the coding and designing of the reality monitoring experiment. HK helped with recruitment of subjects and with designing and acquiring the reality monitoring data. CG helped with initial designing of the reality monitoring experiment. AF helped with analyzing a portion of the reality monitoring MEGI data. JH provided advice on the analyses and interpretation of the MEGI data. SN edited the manuscript, and provided advice on the overall acquisition, analyses and interpretation of all the data.

